# Against the current: upstream behavior in diatoms

**DOI:** 10.64898/2026.02.10.705029

**Authors:** Leonilde Roselli, Giusy Giugliano, Emilie Houliez, Chiara Pennesi, Lisa Miccio, Vittorio Bianco, Pietro Ferraro

**Affiliations:** Stazione Zoologica Anton Dohrn, Brindisi Marine Centre, Department of Research Infrastructures for Marine Biological Resources, Via Duomo 20, 72100 Brindisi, Italy; NBFC, National Biodiversity Future Center, Piazza Marina 61, 90133 Palermo, Italy; CNR-ISASI, Institute of Applied Sciences and Intelligent Systems, Via Campi Flegrei 34, 80078 Pozzuoli, Napoli, Italy; Department of Mathematics and Physics, University of Campania, Viale Abramo Lincoln 5, 81100 Caserta, Italy; Stazione Zoologica Anton Dohrn – CRIMAC, Calabria Marine Centre, Department of Research Infrastructures for Marine Biological Resources, C.da Torre Spaccata, 87071 Amendolara, CS, Italy; Stazione Zoologica Anton Dohrn, Fano Marine Centre, Department of Integrative Marine Ecology, Viale Adriatico 1, 61032 Fano (PU), Italy

## Abstract

Diatoms significantly contribute to aquatic primary productivity and biogeochemical cycles, with motility playing a crucial role in their ecological success. While several factors influence their motility, the effect of water flow remains poorly understood. This study used a digital holographic microscope to investigate the locomotion of the pennate diatom *Navicula* cf *parapontica* under varying flow rates. It demonstrates, for the first time, that *Navicula* perceives and actively counteracts water flows. As flow rates increased up to 500 nL/s, cells consistently moved against the current and frequently adjusted their orientation to maximize resistance. This behaviour allowed the diatoms to maintain a stable locomotion velocity despite a 6.7-fold increase in flow rate. This active rheotaxis likely serves as a strategy to resist resuspension and passive dispersal. These findings reveal a behavioural trait that might play significant role in the way benthic diatom communities maintain their position in the sediments, influencing bentho-pelagic coupling and biogeochemical processes.

## INTRODUCTION

Diatoms play key roles in aquatic ecosystems. They contribute up to 25% of the global primary productivity and drive major biogeochemical cycles^1,2^. Diatoms also have implications for the transfer of energy to higher trophic levels and are crucial in mediating bentho-pelagic coupling^3^. Motility is one of the key traits adopted by diatoms that significantly affect fitness, enabling them to respond to a wide range of environmental factors and exploit spatial heterogeneity. These organisms have evolved for millions of years and developed the ability to actively migrate vertically in aquatic ecosystems. In the pelagic zone diatoms use sinking as a form of motility to explore spatial heterogeneity, adopting several strategies to balance light requirement near the surface and nutrient availability in deeper waters^4-6^. In sedimentary environments, diatoms also have the potential to migrate vertically^7^. In the benthic zone, diatoms employ directed motility on short spatial scales to exploit environmental habitat heterogeneity. The most visible and well described form of motility in benthic diatoms is the vertical migration that occurs daily in the upper sediment layers and is typically synchronized with photoperiod and tidal cycles^8,9^. During daylight low tide (emersion), they move upward to form dense, transient biofilms at the surface of the sediment, optimizing light exposure for photosynthesis. Conversely, they retreat back into deeper sediment layers just before or upon the onset of high tide (immersion) and/or nighttime. This recurrent pattern is continuously adjusted to match the daily and fortnightly tidal cycles, as well as the progressive seasonal changes in day length^7-9^.

Motility is believed to have conferred a critical adaptive advantage to benthic diatoms belonging to the pennates. This advantage would explain their evolutionary success, fast diversification, and why they represent the most recent and most diverse group of diatoms^10,11^. Within the pennates, the raphid diatoms are distinguished by the presence of the raphe, a longitudinal thin and long slit through the valve surface. The raphe is determinant for their ability to achieve directed motility^12^. It has been hypothesized that directed motility favored the biology and ecology of pennate diatoms for a number of reasons. Active motility enables them to enhance nutrient uptake in combination with the capacity to forage using chemotaxis^13^. Motility also facilitates sexual reproduction by enabling the diatoms to reach deep, nutrient-rich layers where reproduction is favored^14^, and helps reduce exposure to grazers by migrating downwards away from the sediment surface where grazers are present^15,16^. Furthermore, vertical migration reduces the negative impacts of sedimentary instability, allowing diatoms to escape from resuspension into the water column^17,18^. It also allows them to anticipate the environmental periodicity, helping cells to avoid main periodic events such as sunrise/sunset, and tidal ebb/flood^8,19^. Finally, motility is linked to the capacity to produce a large amount of mucilaginous extracellular substances (EPS), which provides essential protection against desiccation and high salinities, and serves as a mechanism to deal with photosynthetic overflow^20-22^.

Gliding is the most well-studied type of movement in pennate diatoms because this behavior is easily observable under a microscope^23^. A number of different hypotheses have been proposed to explain this unique mode of cell motility^24,25^. The most widely accepted, proposed by Edgar and Pickett-Heaps^26^, argues for an adhesion/motility complex based on an actin-myosin mechanism. The force for the cell’s back-and-forth movement is generated by the excretion of EPS from its raphe and allows for quasi-instantaneous directional reversal movements towards a stimulus^27,28^. In the absence of a vectorial stimulus, cells tend to move randomly^29^. Several biotic and abiotic factors have been shown to influence and/or trigger diatoms gliding. This includes: light (intensity and spectrum), UV radiation, gravity, chemical gradients, temperature, pH, salinity and calcium^7^. However, there is still a lack of information regarding diatom motility in the presence of water flows. This type of stimulus has not been extensively investigated yet, although it is one of the most predominant cues diatoms are exposed to. Besides, as a result of climate change and temperature rise, one of the expected consequences is the significant variation in the current patterns. Hence, it is important to investigate the effects of such stimuli on benthic diatoms motility and their resiliency. The present study investigated the motility response of the pennate diatom *Navicula* cf *parapontica* when exposed to different flow rates. Species of the genus *Navicula* spp. are frequently reported among the epipelic and epipsammic motile species of microphytobenthic communities. Like many other raphid pennate diatoms, *Navicula* spp. locomotion is based on gliding with excretion of EPS and formation of mucilage trails. Locomotion velocity and trajectories of *Navicula* cf *parapontica* in the presence of different water flow rates were analyzed in a microfluidic laboratory environment. Observations of their locomotion in the presence of water flows were made possible thanks to a self-made digital holographic microscope (Fig. 1). Results show the capability of *Navicula* cf *parapontica* to feel and react to the flow stimulus, and the adaptivity of its response to flow rate increases.

**Figure 1.**
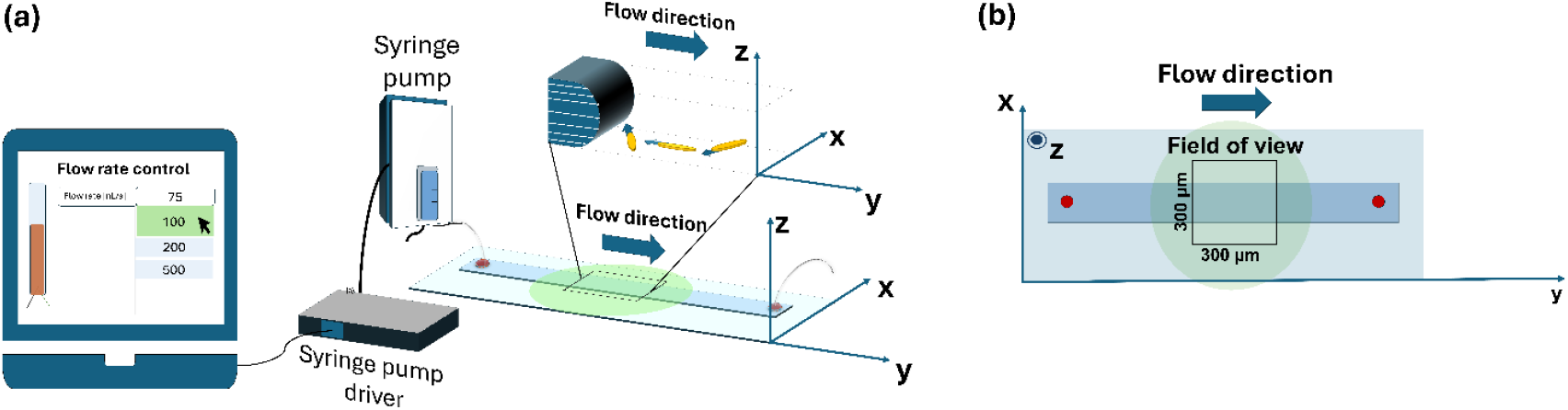
Schematic representation of the in-flow digital holographic microscope setup used to expose Navicula cf parapontica to different flow rates (a). Cell locomotion was tracked in 3D along the x, y and z axes. The blue arrow represents the flow direction along the positive y-axis. (b) Detailed top view of the microfluidic channel, illustrating the field of view and coordinate system for cell tracking.

## RESULTS

### *Navicula* cf *parapontica* motility in presence of an increasing flow rate

In the absence of water flow (control condition), *Navicula* cf *parapontica* cells glided with an apparent random pattern. They moved in all directions without any preference for specific directions (Fig. 2a). Their trajectories were straight or slightly curved with frequent back-and-forth movements and changes in direction. As shown in the pie chart (Fig. 2a), trajectories were distributed between the hemisphere in the opposite direction of the flow (56.1%) and the hemisphere in the same direction as the flow (43.9%) (see Methods section). This near-uniform distribution is further supported by the relative frequency bar plot (Fig. 2a), which shows motion directions balanced across the four sectors A, B, C, and D.

**Fig. 2.**
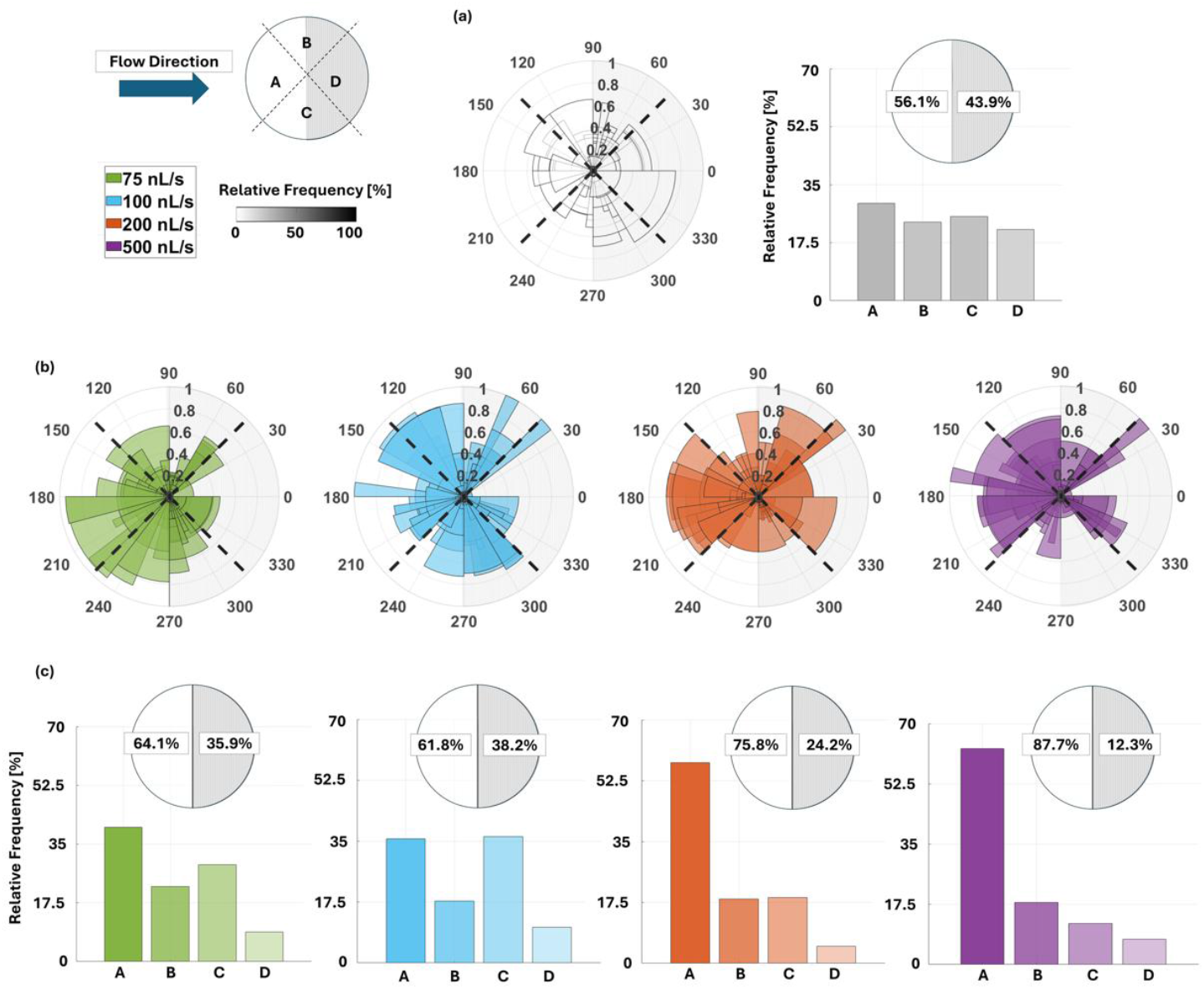
Directions of movement of Navicula cf parapontica cells exposed to increasing flow rates. Polar histograms, relative frequency bar plots and hemisphere division pie charts showing the direction of movement of Navicula cf parapontica cells in the different sectors of the microfluidic channel: a) in the absence of flow and when exposed to a flow rate of b) 75 nL/s, 100 nL/s, 200 nL/s and 500 nL/s. The flow is facing sector A. Cells moving in sector A moved in the opposite direction of the flow, indicated by the blue arrow. Cells moving in sector C moved in the same direction as the flow. Sectors B and C are the cross-flow sectors.

The exposure of cells to a water flow rapidly triggered cell displacement in the direction opposite to the flow (Supplementary Media 1). The pie charts reveal a strong preference in movement direction toward the hemisphere in the opposite direction of the flow, ranging from 61.8% to 87.7% (Fig. 2b). This resulted in the cells moving predominantly into sectors A and C (see relative frequency bar plot in Figs. 2c). As the flow rate increased, the directional preference became more pronounced. The frequency of orientations of the velocity vector into sector C decreased: it was 28.8% and 36.5% at flow rates of 75 and 100 nL/s, respectively, but dropped to 18.9% and 11.9% at 200 and 500 nL/s. Conversely, the frequency of such orientations to sector A increased: it was 40.0% and 35.5% at 75 and 100 nL/s, respectively, but rose to 57.7% and 62.8% at 200 and 500 nL/s.

The cells moved with different trajectories (Fig. 3) but, with increasing flow rates, they tended to favor displacements oblique to the flow direction (e.g. see the zoom-in insets in Fig. 3b).

**Figure 3.**
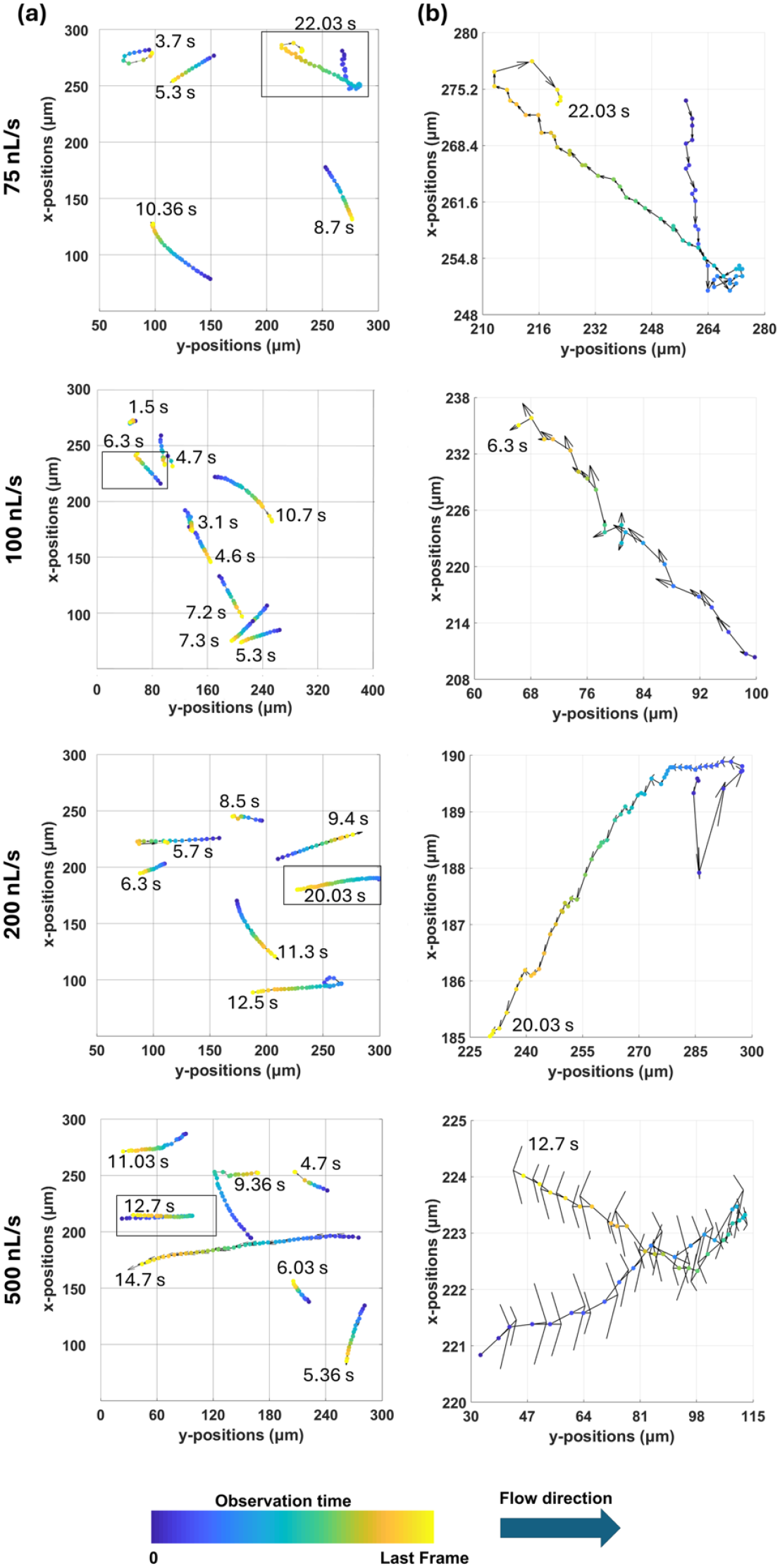
(a) Trajectories of diatoms, color-coded by observation time, exposed to controlled flow rates (75nL/s, 100nL/s, 200nL/s, 500nL/s). (b) Zooming on a single trajectory for each flow rate to show the cells’ micro-movements. Length of vector represents the magnitude of the instantaneous velocity, end extremes of arrows representing the displacement direction that is proportional to the distance travelled between consecutive frames. For each trajectory, the last frame travel time is explicitly stamped with a numerical value indicating the total observation time in seconds (s).

Changes in the direction of movement along the trajectories were frequent at all flow rates, but cells made more frequent orientation changes along their trajectories as the flow rate increased. This trend has been quantified by measuring the standard deviation of the velocity vector orientation vs. the flow rate (Fig. 4a). The relative frequency of changes in orientation increased with increasing flow rates by 3.16% from 75 to 100 nL/s, 11.83% from 100 to 200 nL/s and 31.8% from 200 to 500 nL/s. Cells oriented themselves by exposing the side of their frustule to the incoming flow instead of facing it with their frustule ends. The cells attempted to progress against the flow with the same frustule end forward. In some cases, cells rotated 180° and continued to progress against the flow with the other frustule end forward. With these strategies, the cells were able to maintain a mean velocity of movement that remained relatively stable across the different flow rates tested (Fig. 4b). Despite a nonlinear increase in the input flow rate by a factor of 6.7 (from 75 nL/s to 500 nL/s), the mean velocity of movement measured ranged from 8.9 to 10.9 µm/s, representing a variation by only a factor of 1.2. The capacity of *Navicula* cf *parapontica* to resists water flows obviously presents limits. When the flow rate exceeded the resistance force and locomotion capacity of the cells, the cells were swept away (entrained) and transported by the flow. In the microfluidic channel used, *Navicula* cf *parapontica* cells were unable to resist flow rates higher than 500 nL/s.

**Figure 4.**
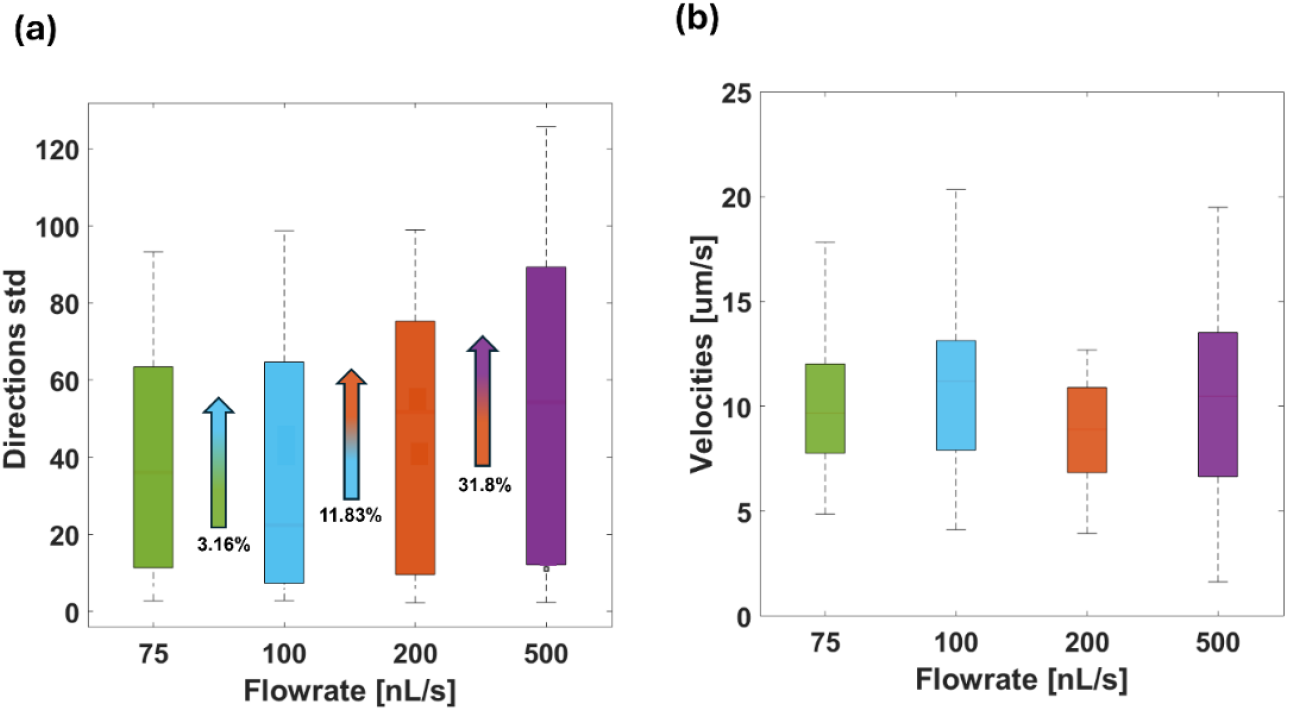
Orientation and velocity of Navicula cf parapontica cells in response to increasing flow rates. (a) Standard deviation of orientation of the velocity vector in response to four flow rates. Arrows represent the relative increase in the frequency of direction changes with the increasing flow rate. (b) Velocity distribution of cells exposed to four different flow rates.

### Short-term response to increasing flow rates at single cell level

The instantaneous velocity of a single *Navicula* cf *parapontica* cell exposed to successive increases in flow rates (Fig. 5) exhibited extreme, rapid fluctuations (ranging from nearly 0 to 30 µm/s) representing periods of accelerations and decelerations consistent with the frequent changes in orientation observed along the gliding trajectory of the cell. However, in spite of these short-term variations in velocity, the mean velocity calculated across the different flow rates fell within a narrow range (8.51 to 14.69 µm/s) indicating that the cell was able to maintain a relatively stable motion velocity. Surprisingly, as the flow rate increased from the moderate (100 nL/s) to the highest flow rates (200 and 500 nL/s), the cell’s mean velocity did not decrease. This behavior suggests the efficiency of the cell’s strategies in generating sufficient propulsive force to successfully counteract the incoming flow. Similarly to what was observed in the first experiment with different individuals exposed to increased flow rates, when a single cell is exposed to successive increasing flow rates, it preferentially moved in the opposite direction of the flow. Furthermore, the cell changed more frequently its orientation along its trajectory as the flow rate increased (Fig. 5b). The standard deviation of the cell orientations along the trajectory equaled 58.30 when exposed to a flow rate of 75 nL/s and progressively increased as the flow rate increased to reach 108.86 at the highest flow rate (500 nL/s). Indeed, the pie charts in Fig. 6b show a strong preference in motion direction toward the hemisphere in the opposite direction of the flow, up to 93.5%. This result is further confirmed by the relative frequency bar plots (Fig. 6b).

**Figure 5.**
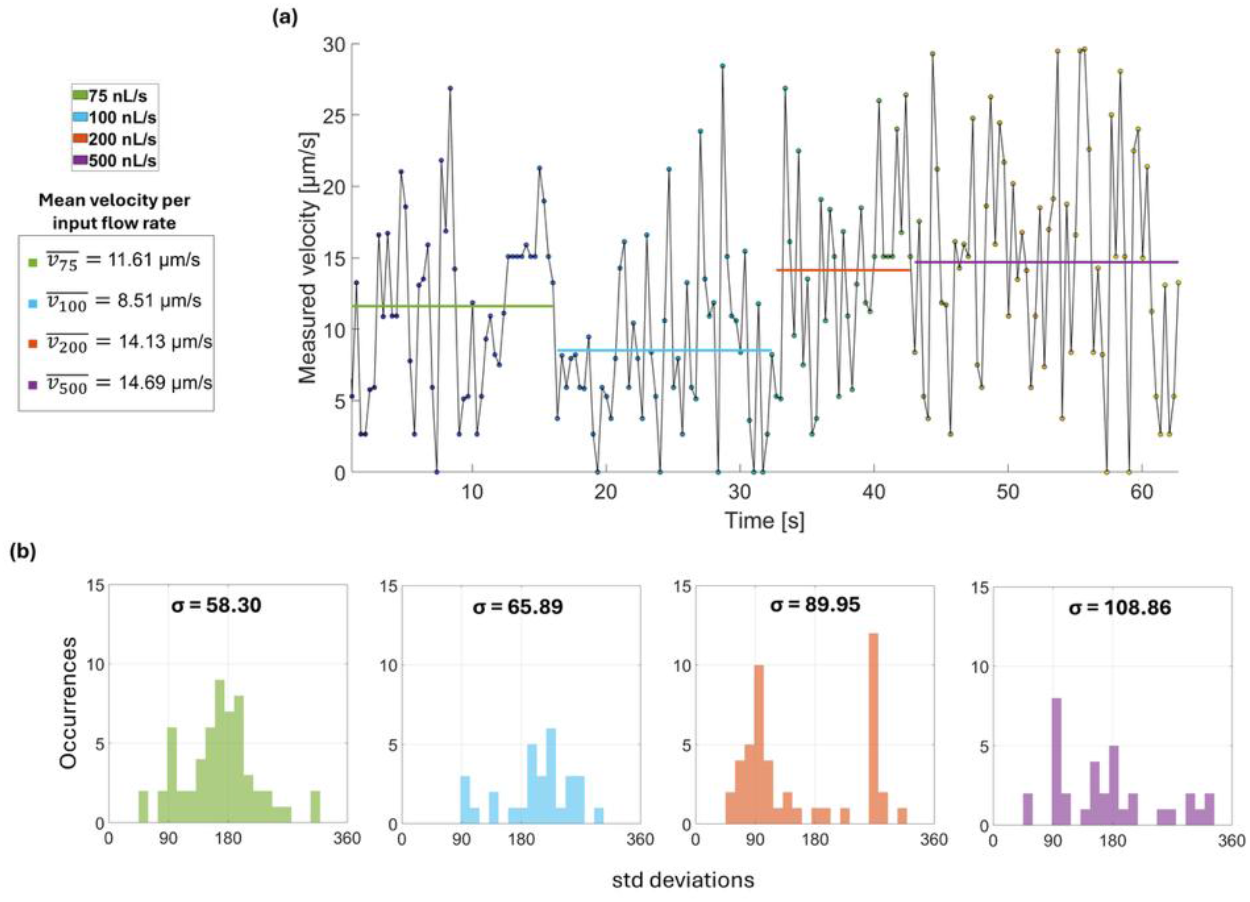
Velocity, directional variability and directionality of one single Navicula probed at different flow rates (75 nL/s, 100 nL/s, 200 nL/s, and 500 nL/s). (a) Measured velocity vs. time. Color-coded shaded regions indicate distinct flow conditions. Mean velocities, 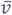, for each flow rate are provided. (b) Histograms of directions of Navicula throughout its trajectory. Standard deviations, σ, for each flow rate reflects the degree of directional variability, showing increased variability with increasing flow rate.

**Figure 6.**
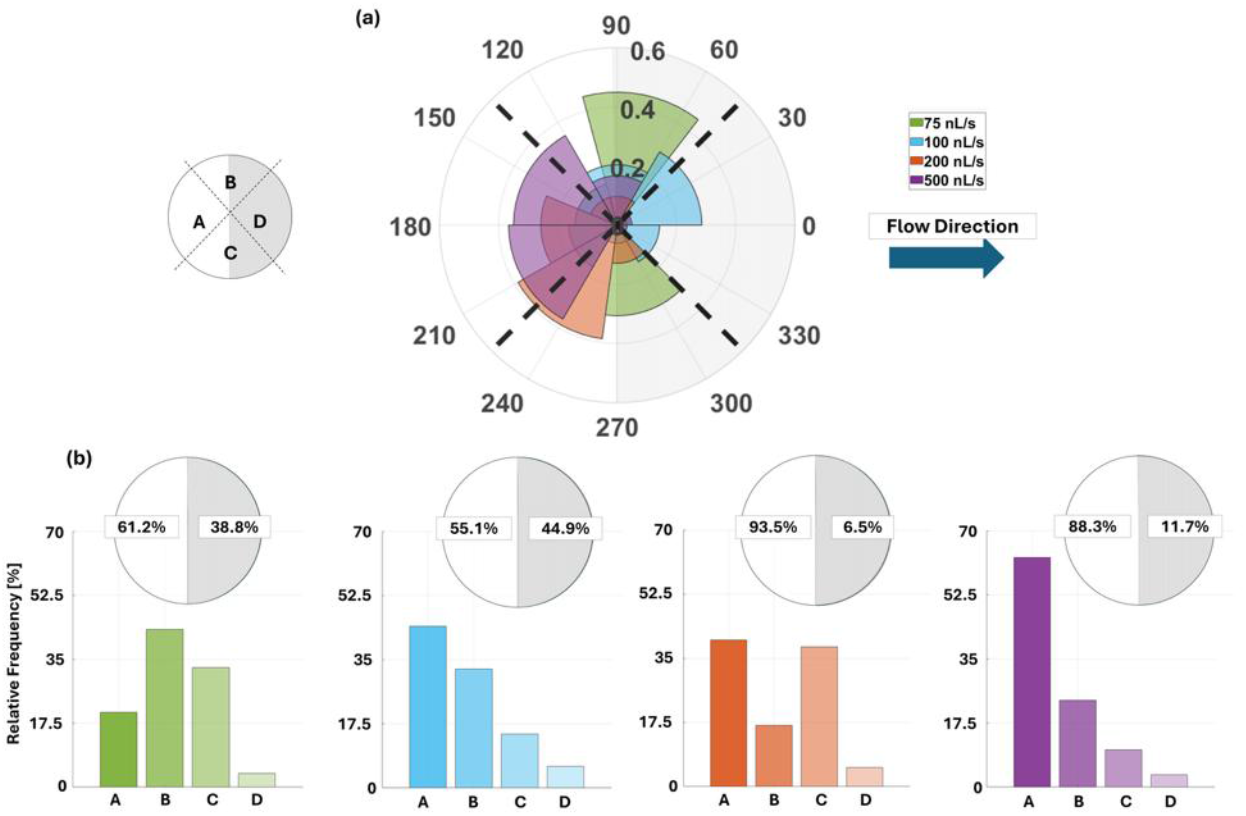
(c) Polar plot summarizing the overall directional distribution across the four flow conditions. (c) Bar plots showing the relative frequency of the cell’s orientation in each of the four angular sectors (A–D) and the pie charts indicate the percentage of trajectories falling within the hemisphere from which the stimulus originates versus the opposite side.

## DISCUSSION

Experimental results clearly demonstrated that *Navicula* cf *parapontica* can perceive water flow direction and speed. A detailed analysis of its movements showed a rapid response to flow rate changes. Indeed, *Navicula* cf *parapontica* tended to move in the opposite direction of the flow to avoid to be swept away. This contrasts with the apparent random locomotion observed in the absence of flow. As flow rates increased, *Navicula* cf *parapontica* more frequently modified its locomotion trajectories while keeping constant average velocity. This behavior was observed at both, the collective and single cell levels. To our knowledge, such locomotion behavior, in response to flow rates, has not been previously described in diatoms.

Although diatom motility has been studied in different species and a large variety of movement modalities^27,30,31^ and gliding speeds^32^ have been reported, none of these studies have documented a diatom locomotion in response to flow rates. This gap may be due to methodological limitations, as diatom motility has mainly been studied in the laboratory using microscopy techniques that do not allow cells to be exposed to water flows. In this study, the development of a self-made digital holographic microscope integrated with a microfluidic platform enabled such experimental designs. This setup allowed for real-time acquisition while the cells were actively exposed to controlled flow conditions, opening new perspectives for the study of diatom motility.

In the field, a study by Passy^33^ suggested that a motile guild of diatoms, including the genus *Navicula*, was more effective at maintaining its relative abundance along a gradient of current velocity than the low profile and high-profile guilds. However, the methodology employed could not clearly demonstrate that their maintenance ability was associated with their motility in response to the current. Results of the present study demonstrated that *Navicula* cf *parapontica* can face water flows. This suggests that in its natural environment, *Navicula* cf *parapontica* is capable of avoiding resuspension during sudden increases in hydrodynamic stress, such as during wind-induced waves or current variations. This capacity to maintain its position in the sediment is likely effective as long as the flow rate does not exceed its locomotion capacity and resistance force. The laboratory experiments showed that, in the microfluidic channels, *Navicula* cf *parapontica* was unable to face a flow rate higher than 500 nL/s. Contrary to phytoplankton that developed the ability to rapidly cope with environmental variations while being transported by the currents, benthic microalgae developed strategies to maintain their position in the sediment and avoid resuspension. They secrete EPS that, in addition to offering them protection against desiccation and high salinity, stabilizes the sediment^8,34^. They also use motility and vertical migrations in anticipation of the incoming tide and/or day/night cycles^8,34,35^. It can be speculated that the capacity to sense and respond to sudden variations in flow rates, as demonstrated in the present study, is an additional strategy. *Navicula* cf *parapontica* cells appeared able to perceive the signal of hydrodynamic changes and instantaneously reacted by “*deciding to stay*”, anchoring to the bottom instead of flowing with the current to seek for a more suitable niche elsewhere. It can be hypothesized that this behavioral trait confers benefits by reducing passive dispersal (due to waves, currents, tides), leading to greater spatial isolation between local populations and increased rates of speciation.

*Navicula* is a large, widespread genus, occurring globally in marine, estuarine, and freshwater environments. Raphe associated motility is considered a recent trait in diatom evolution^36^. It has been hypothesized that direct active motility enables raphid diatoms to colonize new niches and, by enhancing sexual reproduction, to be the primary driver of the rapid and large diversification that has made this group the most diverse of present-day diatoms^7^. The observed motility behavior might be a key mechanism explaining the often, patchy species composition and structure of marine biofilms such as those formed by intertidal microphytobenthos. At the global scale, this has implications by affecting primary productivity and biogeochemical processes.

By using a novel experimental design based on in-flow digital holographic microscopy, this study demonstrated for the first time the capacity of *Navicula* cf *parapontica* to use active locomotion to counter water flows. This suggests that this mechanism might be an additional strategy allowing pennate diatoms to maintain their position in the sediments. The *Navicula* species used in this study was isolated from the Mediterranean Sea, which is a microtidal marine system where pennate diatoms living in the sediment are likely to spend most of their time in submerged conditions. It would be interesting to determine if the *Navicula* species encountered in the intertidal area of macrotidal systems also present this ability. Determining the factors influencing diatom locomotion, growth and functional diversity under ongoing changes in climate and other anthropogenic influences is essential for understanding the functioning of aquatic ecosystems and for predicting their capacity of resilience to changing environmental conditions.

## METHODS

### Isolation and maintenance in culture

*Navicula* cf *parapontica* was isolated from the Gulf of Naples (40°49’N, 14°15’E) located in the Mediterranean Sea. It was maintained in culture in natural filtered seawater from the Gulf of Naples (salinity 36) autoclaved and supplemented with F/2 + Si at a temperature of 22°C and under natural ambient light conditions provided by a window. The species was identified by microscopy by observing the cells using an inverted light microscope Nikon Eclipse Ts2R with a Nikon DC IN camera and a benchtop electron microscope SEM Phenom XL G2 (Supplementary Information, SI. 1).

### Experimental setup

*Navicula* cf *parapontica* locomotion velocity and trajectories were measured using a self-made holographic microscope^37^ (see SI.2 in the supplementary materials for a detailed description of the experimental setup shown in Fig. S3a). Label-free samples of *Navicula* cf *parapontica* were injected into a microfluidic Lab-on-a-Chip device 38,39 and exposed to different flow rates controlled by an automatic syringe pump (CETONI Syringe Pump neMESYS 290N). The experimental assay allowed accurate tracking of each cell. A digital holographic microscope captures in transmission mode holographic video sequences of all the objects moving inside an imaged volume of liquid. This microscopy method is here selected for its capability to refocus and 3D track in post-processing multiple objects acquired severely out-of-focus at the same time^40,46^.

In the first experiment, samples of *Navicula* cf *parapontica* culture were exposed to four different flow rates (75 nL/s, 100 nL/s, 200 nL/s, 500 nL/s). At each flow rate, the locomotion velocity and trajectories of the cells in the field ^1^of view were tracked by recording holograms (see below for more details). The locomotion velocity and trajectories of the cells in the absence of flow were also used as a control. In the second experiment, the locomotion velocity and trajectories of a single cell were tracked by successively exposing it to increasing flow rates (75 nL/s for 16 s, then 100 nL/s for 16 s, 200 nL/s for 10 s and finally 500 nL/s for 20 s). This second experiment was used to analyze the short-term response of *Navicula* cf *parapontica* to variations in flow rate.

### Velocity and trajectories estimation

*Navicula* cf *parapontica* locomotion velocity and trajectories were tracked using frame-by-frame quantitative phase maps (QPMs) extracted from recorded holograms of the entire field of view (see SI.3 in the supplementary materials for QPMs extraction). The motion of each *Navicula* cf *parapontica* cell was characterized by tracking its displacement in three spatial dimensions (x, y, z) over time. The inclusion of the z-dimension (focus depth) allowed for a more comprehensive trajectory description, accounting for the cell’s position relative to the bottom of the microfluidic channel. The tracking was challenging because the *Navicula* cf *parapontica* cells did not consistently flow along the y-axis (parallel to the downstream direction of the flow) nor maintained a constant x-position. To track the cells movement over time, an iterative algorithm was implemented. In the first frame, the diatom’s position was detected, and its centroid was stored as a reference point. For subsequent frames, a binary map was generated, with each identified object corresponding to a distinct moving *Navicula* cf *parapontica* cells. The cell whose centroid exhibited the smallest Euclidean distance relative to the previous reference centroid was recognized as the cell of interest, and its position became the new reference point.

The position of the centroid in the subsequent frames was used to calculate cell’s velocity and characterize its direction of movement. Instantaneous velocity of each cell in x (v_x_) and y (v_y_) directions was obtained by calculating the distance difference between the successive positions of the centroid divided by the corresponding time interval (dt).

### Automatic identification of gliding diatoms

A dual-criteria approach was employed, integrating holographic 3D tracking and observed kinematic behaviour over time to ensure that the analysis was not affected by the inclusion of non-gliding cells. As a preliminary screening criterion, the kinematic direction was assessed to identify cells moving passively with the flow. The kinematic direction was quantified by normalizing the instantaneous velocity vector [v_x_, v_y_] with respect to its total speed, which was computed as the magnitude of the velocity vector in the xy-plane by using the Euclidean norm: 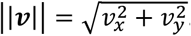. The cosine of the angle between the velocity vector and a reference vector (i.e. the vector describing the flow direction) was calculated using a scalar product. Cells exhibiting a scalar product > 0.9 for more than 20% of their trajectory frames were classified as passive flowing diatoms (for a representative example, see Fig. S3c). These individuals were utilized as tracers to calculate the maximum velocity (i.e.*v*_*max*_), as detailed in SI. 4 of the supplementary materials. The second, more restrictive criterion was established to define the population of cells attached to the microfluidic channel bottom, which is prerequisite for active gliding motility. Adherence was verified by estimating the z-coordinate of the single cell over the trajectory (an example is shown in Fig. S3b, see SI. 3 in the supplementary materials for a detailed description of z-coordinate estimation). A cell was classified as adherent if its calculated z-position remains within ± 1.5 µm of the experimentally determined channel bottom, accounting for the system’s inherent axial resolution (i.e. 1.1 µm), the *Navicula* cf *parapontica* cell thickness, and potential micro-scale surface irregularities.

### Directional analysis and sector partitioning

The kinematic directions of movement of *Navicula* cf *parapontica* were analyzed by constructing polar histograms representing the angle between the velocity vector of each cell and the vector describing the flow direction. For each cell trajectory, instantaneous directions of movement were binned into 30° angular segments delimiting calculated the relative frequency of directions falling within two perfect hemispheres (i.e. [90° - 270°] and [270° - 90°]) to broadly distinguish between downstream and upstream motion. Subsequentially, for a more nuanced analysis and to avoid placing opposite direction of the flow and same direction as the flow (180° and 0°, respectively) directly on bin boundaries, the polar plot was further divided into four 90° quadrants: A, [135° - 225°]; B, [45° - 135°]; C, [225° - 315°]; D, [315° - 45°]. Thus, A is the opposite direction of the flow sector, B and C are the left cross-flow sector and right cross-flow sector, D is the same direction as the flow sector. For each of these four quadrants, we again calculated the relative frequency of directions.

## Acknowledgements

This work was supported by Progetto PRIN 2022 PNRR P2022MA95R-microFluidic platfOrm fOr The toMographicphAse micRoscopy of phytoplanKton assemblages (FOOTMARKS)-CUP B53D23023980001. Funded by European Union-Next Generation EU. This work was partially funded by the National Biodiversity Future Centre (NBFC) Program, Italian Ministry of University and Research, PNRR, Missione 4 Componente 2 Investimento 1.4 (Project: CN00000033). The authors thank Wiebe Kooistra for helping them to isolate *Navicula* cf *parapontica*. The authors acknowledge the use of the SEM provided by the SZN Calabria CRIMAC facility.

## Author contributions

L.R. and V.B. designed the project; V.B., G.G, L.R. and E.H. discussed the experiments; L.M. developed the digital holographic setup; G.G. performed the experiments and cell tracking analysis; L.R., V.B., G.G. and E.H. contributed to interpreting data; C.P. identified the species; E.H. kept the diatom in culture; V.B. and P. F. supervised the work; L.R., G.G and V.B. contributed to drafting the manuscript; all authors contributed to revising the manuscript.

## Additional information

Supplementary Information are included in this paper.

## SI.1. Species description

*Navicula* cf *parapontica* Witkowski, Kulikovskiy, Nevrova & Lange-Bert. In Witkowski et al.^46^, LM description: Frustules rectangular in girdle view. Valves precisely lanceolate with rather acutely rounded, not protracted ends (Fig. S1 a, b).

SEM description: Valves lanceolate with rounded apices 22–37 μm long and 7–11 μm wide. Raphe-sternum is narrow and has a rounded central area. Raphe consists externally of two straight branches ending proximally in expanded central pores and distantly in terminal fissures strongly hooked toward the same side (Fig. S1 f, g, h). The internal raphe branches are straight, ending in a raised central nodule (Fig. S1 c, d) and distally in a raised helictoglossa (Fig. S1 c). Transapical striae (17–20 in 10 μm) parallel over most of the valve and absent near the poles, extending down the mantle. Externally, striae are composed of apically elongated slit-like areolae (Fig. S1 f) that open internally in the same areolae with the same shape (Fig. S1 c).

Our specimens exhibit strong morphological similarities to *Navicula parapontica*^46^; however, we identified some minor differences. *Navicula parapontica* was originally described as a new species by Witkowski et al.^46^ to resolve certain nomenclatural inconsistencies found in previous publications. Specifically, Witkowski et al.^46^ clarified that taxa previously identified as *Navicula pennata* A.W.F. Schmidt var. *pontica* Mereschkowsky sensu Proshkina-Lavrenko^47^, *Navicula pennata* A.W.F. Schmidt var. *pontica* Mereschkowsky sensu Guslyakov et al.^48^, and *Navicula* sp. 134/2 in Witkowski et al.^49^ should all be reassigned to *Navicula parapontica*.The size range of our specimens is comparable in length to *N. parapontica*; however, they tend to be slightly wider (7–11 μm) compared to the 5–6 μm width reported for *N. parapontica*^46^. Additionally, our specimens exhibit strictly parallel transapical striae across the entire valve face (Figs. 1, 2), whereas *N. parapontica* is characterized by transapical striae that are moderately radiate proximally, becoming progressively less radiate to parallel only towards the apices (Fig. 3, p. 311^46^). Furthermore, the number of striae in 10 μm in our specimens is slightly higher (17–20 in 10 μm) than the range reported for *N. parapontica* (12–14 in 10 μm). Finally, both our specimens and *N. parapontica* exhibit slit-like areolae, a key shared feature. However, while *N. parapontica* has a rectangular central area, our specimens display a more rounded central area.

**Fig. S1.**
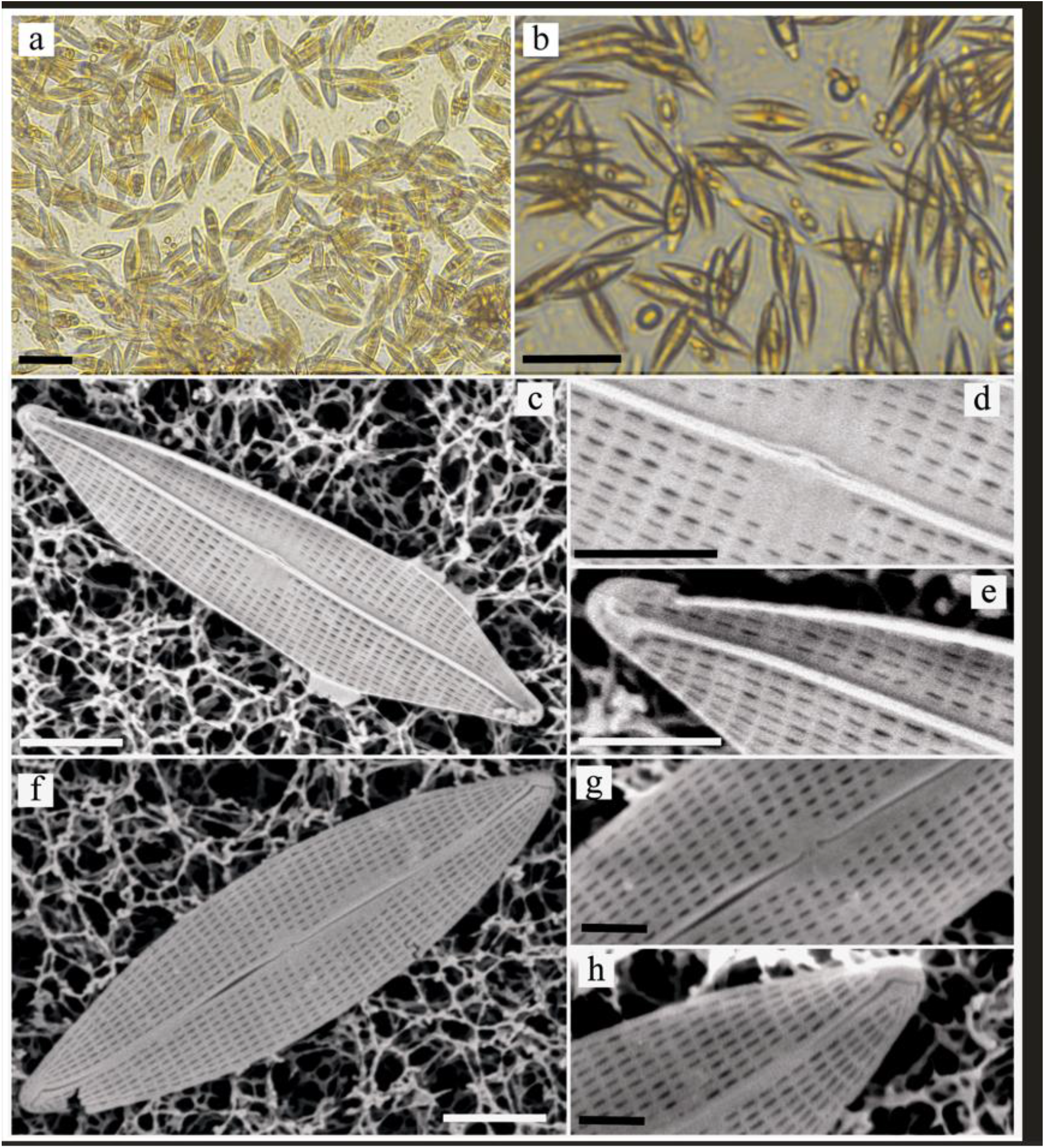
(a-h). *Navicula* cf *parapontica*. LM view. a-b. Scales bars = 30 μm (Figs a-b). Live diatom cells showing the frustule in valve and girdle views. SEM views. c-e. Scales bars = 5 μm (Fig. c), 2 μm (Figs. d, e). Internal valve view showing central nodule (Fig. d) and apical area with helictoglossae (Fig. e). f-h. Scales bars = 5 μm (Fig. f), 1 μm (Figs. g, h). External valve face showing rounded central area (Fig. g) and apical area with deflected terminal fissure (Fig. h).

**Fig. S2.**
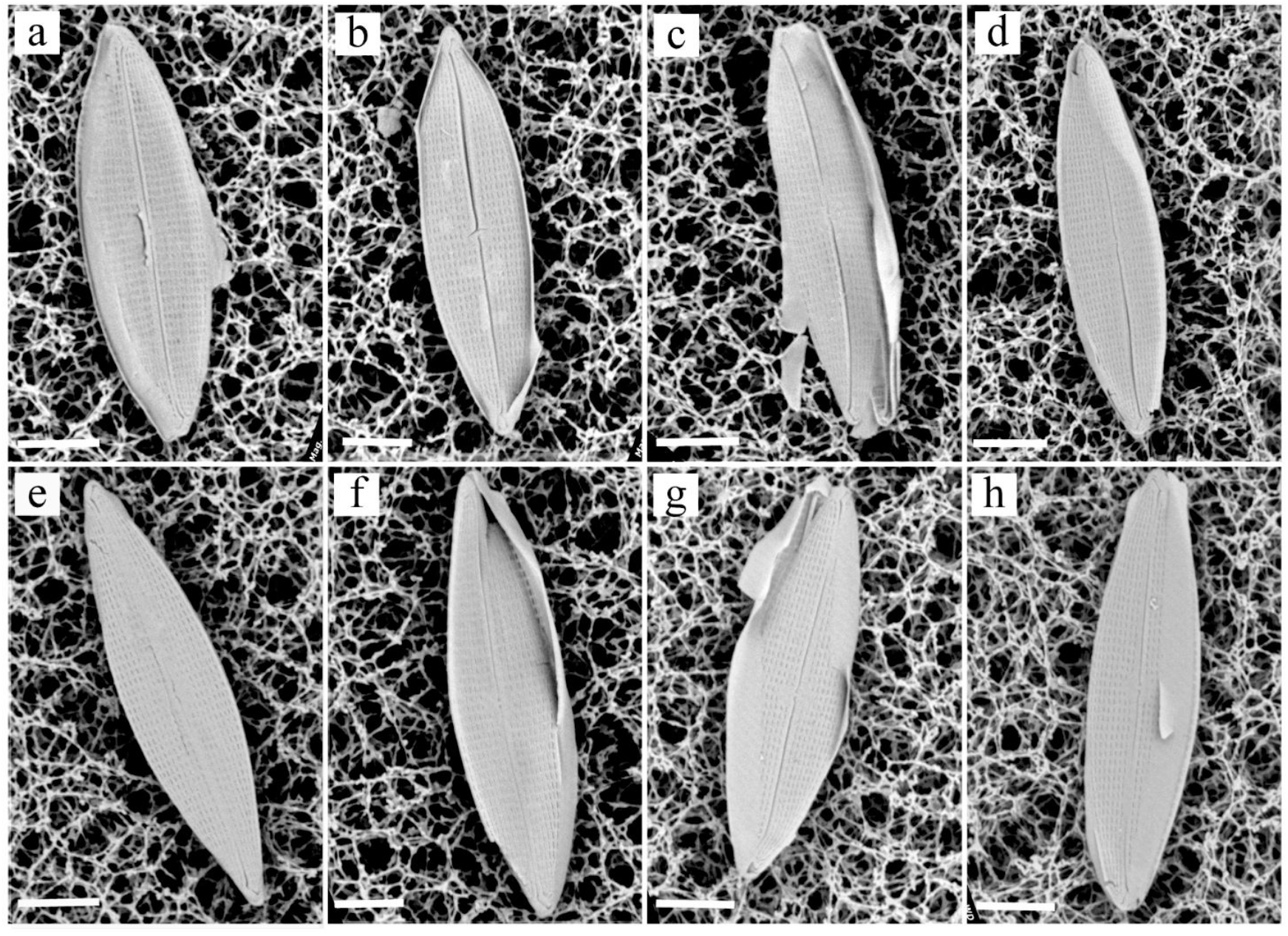
(a-h). *Navicula* cf *parapontica*. SEM view. a-b. Scales bars = 5 μm (Figs a-H). External valve face of different specimens.

## SI.2. Holographic setup

To perform the acquisition of holographic videos (at 30 fps), we implemented a Mach-Zehnder interferometer based on off-axis geometry (Fig. S3a). A waveguided wavelength tunable SuperK laser is set at 495 nm and used as light source. The emitted light wave is collimated and sent into reference and object arms, after the separation operated by a beam splitter (BS1). The object beam passes through a microfluidic channel (MC) having a cross section of 200 µm × 1000 µm and a length of 58.5 mm. The transmitted wavefront is modulated by any object inside the channel and falling within the field of view (FoV). This is collected by a microscope objective (MO1) with a numerical aperture (NA) equal to 0.95, which introduces a 63X sample magnification. The investigated sample is imaged into the primary image plane by a tube lens L1 (achromatic doublet, f = 150 mm) and further re-imaged in by a telescope shaped by two achromatic doublet lenses L1 and L5, with a focal length of 150 mm and 200 mm, respectively. Components with the same paraxial optical parameters are included in the reference arm, adding a diffraction grating G (Newport, Moire Grating 80 lp/mm) in the primary image plane (i.e. between L3 and L4); while, an iris diaphragm (I) is mounted in the Fourier plane of lens L4 for ensuring the exclusive transmission of the first diffraction order. Two mirrors are added to each arm (M1 and M2 in the object arm, M6 and M7 in the reference arm) in order to enhance the temporal stability. The two contributions are re-combined by a second beam splitter (BS2) and the resulting interference pattern is collected by the CMOS camera (Genie Nano-CXP Camera, 5120 x 5120 pixels, Δx = Δy = 4.5 μm pixel size). Since the overall lateral magnification at the camera plane reaches 76.4, the recorded FoV measures about 301 x 301 μm^2^. Theoretically, the maximum spatial frequency that can be transmitted through the imaging system is 1919 lp/mm that corresponds to a lateral resolution of about 0.5 µm. The *Navicula* diatoms were injected into the MC by employing an automatic syringe pump (CETONI Syringe Pump neMESYS 290N) that allows controlling the flow rate.

**Figure S3.**
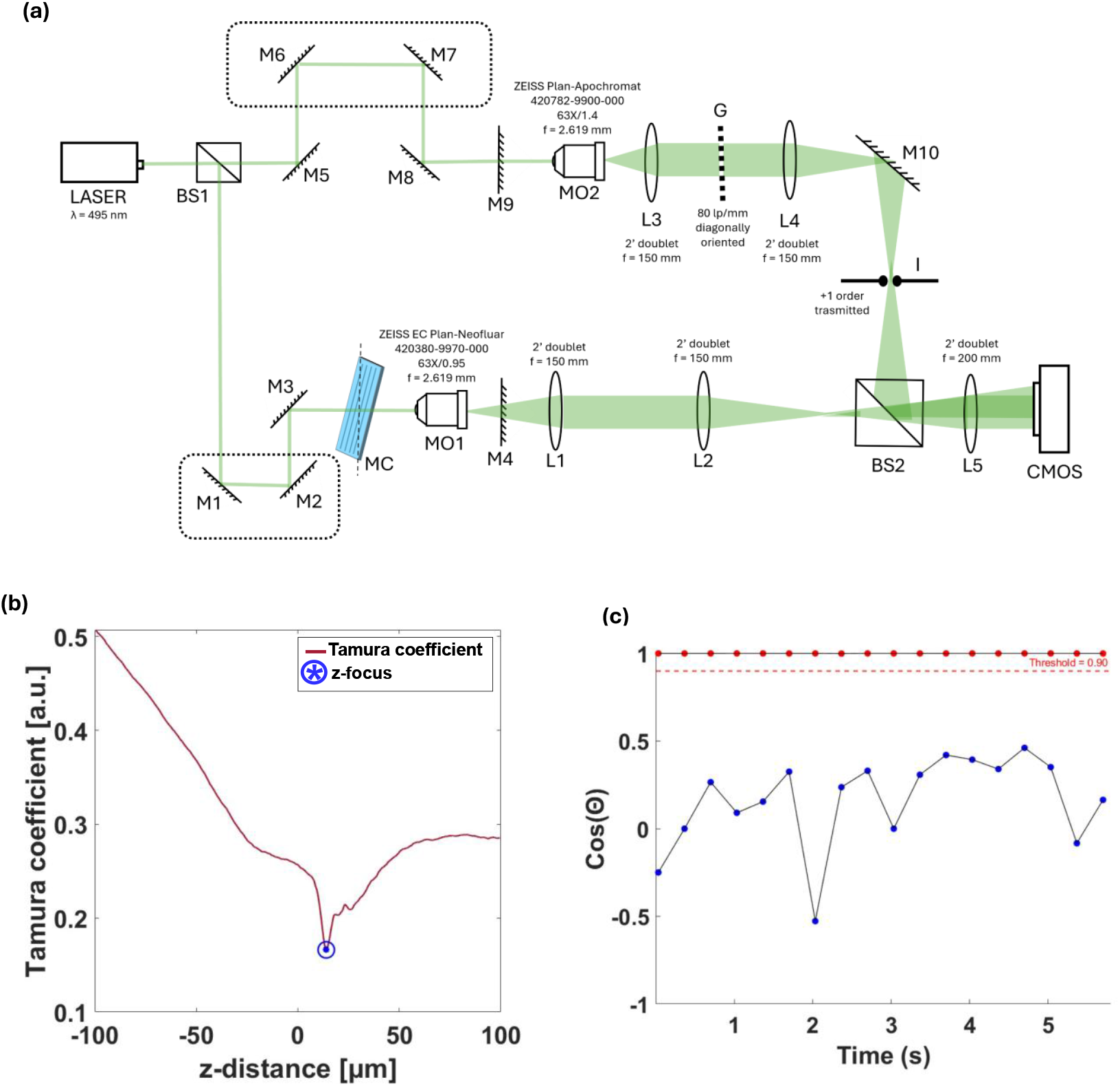
Optical setup and computational analysis for diatom tracking and numerical refocusing. (a) Schematic sketch of the digital holography microscopy setup. The setup includes a laser source (λ = 495 nm), beam splitters (BS1, BS2), multiple mirrors (M1–M10), lenses (L1–L5), microscopy objectives (MO1, MO2), and a CMOS camera for hologram recording. (b) Tamura coefficient as a function of z-distance, used for numerical refocusing. The blue marker highlights the determined in-focus z-position of the tracked Navicula diatom. (c) Temporal evaluation of the cosine of the direction angle (cos(Θ)) of a tracked diatom, with a threshold line (red) indicating the straight downstream direction.

## SI.3. Holographic flow cytometer setup

The interference between object and reference beams allows accessing the complex amplitude of the sample, i.e. both the phase-contrast distribution and the amplitude of the optical field modulated by the object and collected by the camera. The phase-contrast information is encoded by the interference process into a recordable intensity image, namely the hologram, which is the raw camera readout. After numerical reconstruction of the hologram [S1] we achieved the QPMs, frame-by-frame, of the full FoV in which *Navicula* diatoms were imaged while moving. Our approach leverages the *a posteriori* focusing capability of digital holography to automatically differentiate *Navicula* diatoms that adhere to the channel bottom from those that are freely flowing. First, we demodulate the hologram by applying a pupil filtering in the Fourier domain, which allows avoiding the unwanted diffraction orders (i.e. the -1 and zero-th orders of diffraction, which represent the virtual image and the DC term, respectively). Then, the demodulated hologram is back-propagated by applying the angular spectrum propagation method to solve numerically the Rayleigh-Sommerfeld diffraction integral under the assumptions of the scalar diffraction theory [S2]. Then, we estimate the best-focus distance by minimizing the Tamura coefficient (TC) [S3] and propagate the demodulated hologram to such distance, achieving the in-focus complex amplitude, from which the QPM is extracted. In order to get rid of the optical aberrations of the system, we acquire and subtract a reference hologram [S4], that is a hologram acquired without the sample in the FoV. Once the aberrations are properly compensated, we apply a phase unwrapping algorithm based on the least-square approach [S5] that minimizes the error between the observed wrapped phase and the unwrapped phase [S6]. The last step is the denoising of the unwrapped and compensated phase image, by applying the two-dimensional windowed Fourier transform filtering [S7]. The achieved QPM of each FoV was used for tracking the trajectories of the moving *Navicula* diatoms.

In order to obtain regions of interest (ROIs) from the full-FoV hologram, the collected centroids were used to superimpose a centered patch of 700 × 700 pixels containing the detected *Navicula* diatom. On each sub-hologram we applied the numerical reconstruction pipeline that is described above for achieving the proper in-focus distance for each diatom, which provides the axial position of each sample (Fig. S3b). The z position of each sample at each frame is obtained using the digital holography refocusing process described above. Since the TC is described by a convex functional, the proper in-focus distance is defined as the z which corresponds to the global minimum of the TC trend.

## SI.4. Estimation of the parabolic velocity profile

In the MC, a laminar flow is established, resulting in a parabolic velocity profile that can be described using the Navier-Stokes equations [S8] in vectorial form:

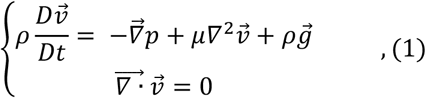

where *ρ* is the density of the fluid, 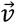 is the velocity of the fluid, *µ* is its viscosity, *p* is the exerted pressure, and 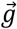 is the gravity force. Solving the Navier-Stokes equations is not a trivial task [S8] due to the non-linearity of the convection terms 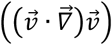. However, in some cases non-linear terms can be neglected, thus making the determination of the exact analytical solutions possible.

**Figure S4.**
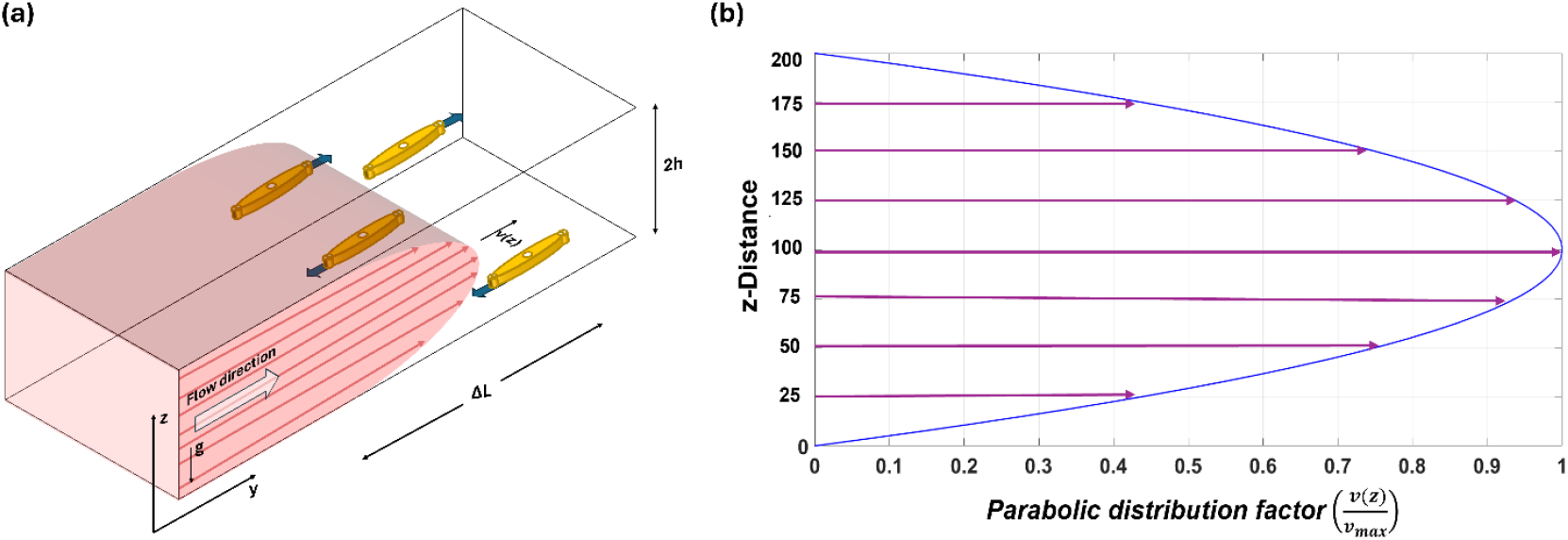
Schematic representation of diatom motion in a microfluidic channel and corresponding parabolic flow profile. (a) Illustration of Navicula diatoms moving within a microfluidic channel under laminar flow condition. The velocity profile follows a parabolic distribution, with maximum velocity v_max_at the center and null velocity at the channel walls. (b) Analytical representation of the parabolic velocity distribution along the z-axis, normalized by v_max_.

A fluid flows between two parallel plates that are distant 2*h* from each other, with the sole non-null component along the y-axis. Considering the fixed refence system sketched in Fig. S4a and the incompressibility of the fluid [S9]., the mass conservation equation can be written as:

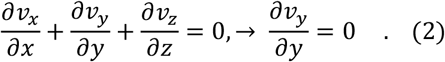

Therefore, by adding the hypothesis of stationarity, the Navier-Stokes equations are reduced to:

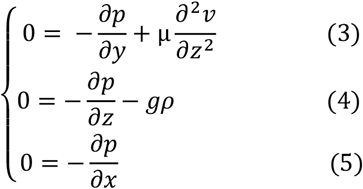

From Eq. (5), it is evident that the exerted pressure,*p*, remains constant across the x direction. Thus, it can be expressed as *p* = *p*(*y, z*). By integrating Eq. (4) with respect to *z*, the exerted pressure, *p*, can be expressed as:

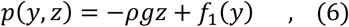

where *f*_1_(*y*) is an arbitrary function that describes the pressure distribution along the y axis. Thus, by fixing y, the exerted pressure varies hydrostatically along the z axis. By differentiating Eq (6) with respect to y, the partial derivative 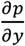 is simplified to 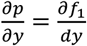. Therefore, the resulting equation is obtained by replacing this result into Eq. (3):

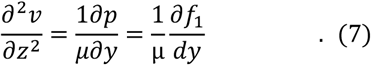

The velocity *v* of the fluid can be obtained by integrating Eq. (7) with respect to *z* as follows:

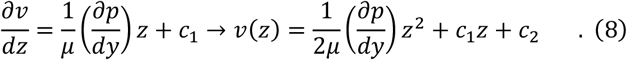

It is important to note that the term 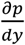 is treated as a constant during the integration in the z direction, which follows directly from the form of Eq. (6).

The constants *c*_1_and *c*_2_ may be determinated by introducing the boundary conditions [S10] *v* = 0|_*z*=±*h*_:

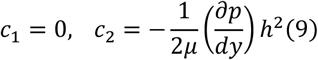

Thus, the final equation of the velocity profile is obtained as follows:

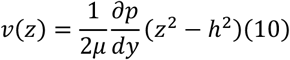

The mathematical representation of the velocity profile follows a quadratic function with the *z* coordinate, indicating that the velocity reaches its peak at the center of the channel (z=0 in the chosen reference system) and gradually reduces to zero at the walls [S11,S12]. In order to avoid the dependence on the pressure gradient, fluid viscosity and mean velocity, it is useful to express Eq. (10) in dimensionless form by normalizing it with respect to the maximum velocity *v*_*max*_, achieving the parabolic distribution factor:

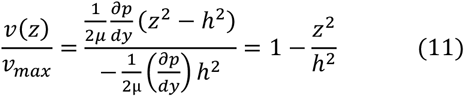

Eq. (11) was involved in the evaluation of the parabolic distribution factor at each distance z (Fig. S4b) using the dimensional parameters of the employed MC (i.e. *h* = 200 µm). Once we identified the flowing diatoms, using the criterion explained in the Methods section, their z-position and velocity distribution were estimated for each flow rate by minimizing the TC and applying the Euclidean norm, respectively. Both z-positions and velocities are reported as z= µ ±3σ for each distribution across all the input flowrates, where µ is the mean of the distribution and 3σ is the uncertainty interval. For the recognized flowing diatoms, we measured z-distance of 19.33 ± 3.35µm, 17.86 ± 3.46µm, 18.45 ± 2.94µm and 23.68 ± 2.13 µm for flow rates of 75nL/s, 100nL/s, 200nL/s and 500 nL/s, respectively. Correspondingly, velocities were recorded at 42.37 ± 14.25µm/s, 51.52 ± 14.82µm/s, 91.12 ± 21.31µm/s and 246.01 ± 59.79 µm/s. By applying the Eq. (11), the *v*_*max*_, i.e. the maximum velocity of the flow in the center of the microfluidic channel, is 121.32 µm/s at 75 nL/s, 158 µm/s at 100 nL/s, 272.03 µm/s at 200 nL/s and 589.20µm/s at 500 nL/s.

## References

1. Field, C. B., Behrenfeld, M.J., Randerson J.T. Falkowski, P. Primary production of the biosphere: integrating terrestrial and oceanic components. Science 281, 237–240 (1998).

2. Smetacek, V. Diatoms and the ocean carbon cycle. Protist 150, 25–32 (1999).

3. Yool, A. & Tyrrell, T. Role of diatoms in regulating the ocean’s silicon cycle. Global Biogeochem. Cycles 17, 1103 (2003).

4. Margalef, R., Life-forms of phytoplankton as survival alternatives in an unstable environment. Oceanol. Acta 134, 493–509 (1978).

5. Gemmell, B.J., Oh, G., Buskey, E.J., Villareal, T.A. Dynamic sinking behavior in marine phytoplankton: rapid changes in buoyancy may aid in nutrient uptake. Proc. R. Soc. B 283, 20161126 (2016).

6. Durante, G., Basset, A., Stanca, E., Roselli, L. Allometric scaling and morphological variation in sinking rate of phytoplankton. J. Phycol. 55, 1386–1393 (2019).

7. Serôdio, J. Diatom Motility: Mechanisms, Control and Adaptive Value. In Diatom Gliding Motility (eds Cohn, S., Manoylov, K. & Gordon, R.) 159–184 (2021).

8. Consalvey, M., Paterson, D. M. & Underwood, G. J. C. The ups and downs of life in a benthic biofilm: migration of benthic diatoms. Diatom Res. 19, 181–202 (2004).

9. Barnett, A., Méléder, V., Dupuy, C., and Lavaud, J. The Vertical Migratory Rhythm of Intertidal Microphytobenthos in Sediment Depends on the Light Photoperiod,Intensity, and Spectrum: Evidence for a Positive Effect of Blue Wavelengths. Front. Mar. Sci. 7:212 (2020).

10. Cahoon, L. B. in Oceanography and Marine Biology: An Annual Review Vol. 37 (eds Ansell, A. D., Gibson, R.N. & Barnes, M.) 47–86 (University College London Press, 1999).

11. Bowler, C., Allen, A., Badger, J. et al. The Phaeodactylum genome reveals the evolutionary history of diatom genomes. Nature 456, 239–244 (2008).

12. Nakov, T., Beaulieu, J.M., Alverson, A.J. Accelerated diversification is related to life history and locomotion in a hyperdiverse lineage of microbial eukaryotes (Diatoms, Bacillariophyta). New Phytol. 219, 462 – 473 (2018).

13. Bondoc, K.G.V., Heuschele, J., Gillard, J., Vyverman, W., Pohnert, G. Selective silicate-directed motility in diatoms. Nat. Commun. 7: 10540 (2016).

14. Saburova, M.A. & Polikarpov, I.G. Diatom activity within soft sediments: behavioural and physiological processes. Mar. Ecol. Prog. Ser. 251, 115 – 126 (2003).

15. Joint, I., Gee, J., Warwick, R. Determination of fine-scale vertical distribution of microbes and meiofauna in an intertidal sediment. Marine Biol. 72, 157 – 164 (1982).

16. Orvain, F. & Sauriau, P.G. Environmental and behavioural factors affecting activity in the intertidal gastropod Hydrobia ulvae. J. Exp. Mar. Biol. Ecol. 272, 191 – 216 (2002).

17. Heckman, C.W. The development of vertical migration patterns in the sediments of estuaries as a strategy for algae to resist drift with tidal currents. Int. Rev. Gesamten Hydrobiol. Hydrogr. 70, 151 – 164 (1985).

18. Kingsto, M.B. Wave effects on the vertical migration of two benthic microalgae: Hantzschia virgata var. intermedia and Euglena proxima. Estuaries 22, 81–91 (1999).

19. Coelho, H., Vieira, S., Serôdio, J. Endogenous versus environmental control of vertical migration by intertidal benthic microalgae. Eur. J. Phycol. 46, 271 – 281 (2011).

20. Abdullahi, A.S., Underwood, G.J.C., Gretz, M.R. Extracellular matrix assembly in diatoms (Bacillariophyceae). V. Environmental effects on polysaccharide synthesis in the model diatom, Phaeodactylum tricornutum. J. Phycol. 42, 363–378 (2006).

21. Perkins, R.G., Lavaud, J., Serôdio, J., Mouget, J.L., Cartaxana, P., Rosa, P., Jesus, B.M. Vertical cell movement is a primary response of intertidal benthic biofilms to increasing light dose. Mar. Ecol. Prog. Ser. 416, 93 – 103 (2010).

22. Steele, D.J., Franklin, D.J., Underwood, G.J.C. Protection of cells from salinity stress by extracellular polymeric substances in diatom biofilms . Biofouling 30, 987 – 998 (2014).

23. Lauterborn, R. Untersuchungen über Bau, Kernteilung und Bewegung der Diatomeen, Leipzig: W. Englemann. 165 pp (1896)

24. Wang, J., Cao, S., Du., Chen, D. Underwater locomotion strategy by a benthic pennate diatom Navicula sp. Protoplasma 250, 1203–1212 (2013).

25. Poulsen, N., Davutoglu, M.G., Zackova Suchanova, J. Diatom Adhesion and Motility in The Molecular Life of Diatoms (eds Falciatore, A., Mock, T.) 367–387 (Springer Nature Switzerland, 2022).

26. Edgar, L. A. & Pickett-heaps, J. D. Diatom Locomotion in Progress in Phycological Research (eds. Round, F.E. & Chapman, D.J.) 3 47–88 (Biopress Ltd, Bristol, 1984).

27. Apoya-Horton, M.D., Yin, L., Underwood, G.J.C., Gretz, M.R. Movement modalities and responses to environmental changes of the mudflat diatom Cylindrotheca closterium (Bacillariophyceae). J. Phycol. 42, 379 – 390 (2006).

28. Cohn, S.A., Bahena, M., Davis, J.T., Ragland, R.L., Rauschenberg, C.D., Smith, B.J., Characterisation of the diatom photophobic response to high irradiance. Diatom Res. 19, 167 – 179 (2004).

29. McLachlan, D.H., Brownlee, C., Taylor, A.R., Geider, R.J., Underwood, G.J.C. Light-induced motile responses of the estuarine benthic diatoms Navicula perminuta and Cylindrotheca closterium (Bacillariophyceae). J. Phycol. 45, 592 – 599 (2009).

30. Cohn, S.A., Farrell, J.F., Munro, J.D., Ragland, R.L., Weitzell, R.E., Wibisono, B.L. The effect of temperature and mixed species composition on diatom motility and adhesion. Diatom Res. 18, 225 – 243 (2003).

31. Jönsson, B., Sundbäck, K., Nilson, C. An upright life-form of an epipelic motile diatom: on the behaviour of Gyrosigma balticum. Eur. J. Phycol. 29, 11 – 15 (1994).

32. Cohn, S.A. & Disparti, N.C. Environmental factors influencing diatom cell motility. J. Phycol. 30, 818 – 828 (1994).

33. Passy, S.I. Diatom ecological guilds display distinct and predictable behavior along nutrient and disturbance gradients in running waters. Aquat. Bot. 86, 171–178 (2007)

34. Serôdio, J., Ezequiel, J., Barnett, A., Mouget, J., Méléder, V., Laviale, M., Lavaud, J. Efficiency of photoprotection in microphytobenthos: role of vertical migration and the xanthophyll cycle against photoinhibition. Aquat. Microb. Ecol. 67, 161 – 175 (2012).

35. Ní Longphuirt, S., Leynaert, A., Guarini, J.M., Chauvaud, L., Claquin, P., Herlory, O., Amice, E., Huonnic, P., Ragueneau, O. Discovery of microphytobenthos migration in the subtidal zone. Mar. Ecol. Prog. Ser. 328, 143–154 (2006).

36. Bowler, C., Vardi, A., Allen, A.E. Oceanographic and biogeochemical insights from diatom genomes. Annu. Rev. Mar. Sci. 2, 333 – 365 (2010).

37. Borrelli, F., Giugliano, G., Houliez, E., Behal, J., Pirone, D., Roselli, L., Sardo, A., Zupo, V., Costantini, M., Miccio, L., Memmolo, P., Bianco, V., Ferraro, P. 3D Holographic Flow Cytometry Measurements of Microalgae: Strategies for Angle Recovery in Complex Rotation Patterns. Lab Chip, 25, 5283–5291 (2025).

38. Leung, C.M., de Haan, P., Ronaldson-Bouchard, K. et al. A guide to the organ-on-a-chip. Nat Rev Methods Primers 2, 33 (2022).

39. Tanriverdi, S., Cruz, J., Habibi, S. et al. Elasto-inertial focusing and particle migration in high aspect ratio microchannels for high-throughput separation. Microsyst Nanoeng 10, 87 (2024).

40. Memmolo, P., Miccio, L., Paturzo, M., Caprio, G. D., Coppola, G., Netti, P. A., & Ferraro, P. Recent advances in holographic 3D particle tracking. Adv. Opt. Photonics 7, 713–755 (2015).

41. Wang, Z., Giugliano, G., Behal, J., Schiavo, M., Memmolo, P., Miccio, L., Grilli, S., Nazzaro, F., Ferraro, P., Bianco, V. All-optical dual module platform for motility-based functional scrutiny of microencapsulated probiotic bacteria. Biomed. Opt. Express 15, 2202–2223 (2024).

42. Wagner, O., Edri, E., Hadikahani, P., Shpaisman, H., Zalevsky, Z., and Psaltis, D. Microfluidic based linear-optics label-free imager. Lab Chip, 1259–1266 (2020).

43. Kim, J., Lee, S.J. Digital in-line holographic microscopy for label-free identification and tracking of biological cells. Mil. Med. Res. 11, 38 (2024).

44. Stollmann, A., Garcia-Guirado, J., Hong, JS. et al. Molecular fingerprinting of biological nanoparticles with a label-free optofluidic platform. Nat. Commun. 15, 4109 (2024).

45. Wang, Z., Bianco, V., Maffettone, P.L., Ferraro P. Holographic flow scanning cytometry overcomes depth of focus limits and smartly adapts to microfluidic speed. Lab on a Chip 12, (2023).

46. Witkowski A., Kulikovskiy M., Nevrova E., Lange-Bertalot H., Gogorev R. The genus Navicula in ancient basins. I. Two novelties from the Black Sea. Pl. Ecol. Evol. 143, 307–317 (2010).

47. Proshkina-Lavrenko A.I. Benthic diatoms of the Black Sea. AS USSR Moscow-Leningrad, Russia, 243 pp. (1963).

48. Guslyakov N., Zakordonez O., Gerasimuk V. Atlas of benthic diatoms of the Northwestern part of the Black Sea and adjacent regions. Naukova Dumka, Kiev., Russia 115 pp. (1992).

49. Witkowski A., Lange-Bertalot H., Metzeltin D. Diatom Flora of Marine Coasts 1. Iconographia Diatomologica 7, 1–926 (2000).

## References

S1. Cotte Y., Toy F., Jourdain P., Pavillon N., Boss D., Magistretti P., Marquet P. and Depeursinge C. Nat. Photonics 7, 113–117 (2013)

S2. Schnars U. and Jüptner W. Meas. Sci. Technol. 13, R85–R101 (2002)

S3. Memmolo P., Distante C., Paturzo M., Finizio A., Ferraro P. and Javidi B. Opt. Lett. 36, 1945– 1947 (2011).

S4. Zhou W., Yu Y. and Asundi A. Opt. Lasers Eng. 47, 264–270 (2009).

S5. Guo Y., Chen X. and Zhang T. Opt. Lasers Eng. 63, 25–29 (2014).

S6. Bioucas-Dias J. M. and Valadao G. IEEE Trans. Image Process. 16, 698–709 (2007).

S7. Qian K. Appl. Opt. 43, 2695–2702 (2004).

S8. Łukaszewicz L., Kalita P. and Grzegorz G. Adv. Mech. Math. 34, 24–25 (2016).

S9. Liron N. and Mochon S. J. Eng. Math. 10, 287–303 (1976).

S10. Amrouche C. and Rejaiba A. J. Differ. Equations 214, 1515–1547 (2014).

S11. Giorgini A., Miranville A. and Temam R. SIAM J. Math. Anal. 51, 2535–2574 (2019).

S12. Foias C., Rosa R. M. and Temam R. J. Dyn. Differ. Equations 31, 1689–1741 (2019).

